# Ribosome quality control activity potentiates vaccinia virus protein synthesis during infection

**DOI:** 10.1101/2020.11.12.380634

**Authors:** Elayanambi Sundaramoorthy, Andrew P. Ryan, Amit Fulzele, Marilyn Leonard, Matthew D. Daugherty, Eric J. Bennett

**Affiliations:** Section of Cell and Developmental Biology, Division of Biological Sciences, University of California, San Diego, La Jolla, CA 92093, USA; Section of Molecular Biology, Division of Biological Sciences, University of California, San Diego, La Jolla, CA 92093, USA

## Abstract

Ribosomes are highly abundant cellular machines that perform the essential task of translating the genetic code into proteins. Cellular translation activity is finely tuned and proteostasis insults, such as those incurred upon viral infection, activate stress signaling pathways that result in translation reprogramming. Viral infection selectively shuts down host mRNA while redistributing ribosomes for selective translation of viral mRNAs. The intricacies of this selective ribosome shuffle from host to viral mRNAs are poorly understood. Here, we uncover a role for the ribosome associated quality control (RQC) factor ZNF598, a sensor for collided ribosomes, as a critical factor for vaccinia virus mRNA translation. Collided ribosomes are sensed by ZNF598, which ubiquitylates 40S subunit proteins uS10 and eS10 and thereby initiates RQC-dependent nascent chain degradation and ribosome recycling. We show that vaccinia infection in human cells enhances uS10 ubiquitylation indicating an increased burden on RQC pathways during viral propagation. Consistent with an increased RQC demand, we demonstrate that vaccinia virus replication is impaired in cells which either lack ZNF598 or contain a ubiquitylation deficient version of uS10. Using SILAC-based proteomics and concurrent RNAseq analysis, we determine that host translation of vaccinia virus mRNAs is compromised in cells that lack RQC activity as compared to control cells whereas there was little evidence of differences in host or viral transcription. Additionally, vaccinia virus infection resulted in a loss of cellular RQC activity, suggesting that ribosomes engaged in viral protein production recruit ZNF598 away from its function in host translation. Thus, co-option of ZNF598 by vaccinia virus plays a critical role in translational reprogramming that is needed for optimal viral propagation.

## Introduction

The task of translating the genetic code into functional proteins is an essential and resource intensive process(Warner 1999). Dividing cells need to double their proteome content prior to cell division to maintain proteome complexity after division. Ribosomes and the translation associated machinery can be limiting for this process and studies in single cell systems have demonstrated that ribosome content sets cellular proliferation rate(Warner 1999; Scott et al. 2010; Scott and Hwa 2011; Scott et al. 2014; Kafri et al. 2016). Interventions that reduce translation speed constrain cell growth and proliferation(Scott and Hwa 2011). Given these observations, it is not surprising that one of the most immediate cellular responses to proteotoxic stress is to reduce protein biogenesis output(Costa-Mattioli and Walter 2020). This reduction in translation has two beneficial outcomes. First, it reduces protein output to limit the production of misfolded, mistranslated, or otherwise faulty proteins that require quality control-dependent degradation. Second, limiting translation allows for resource reallocation to mount a successful stress response to restore cellular homeostasis(Costa-Mattioli and Walter 2020). Viral infection results in proteotstasis imbalance as the virus rewires the cellular translation machinery to rapidly generate new viral particles(Walsh and Mohr 2011).Viral induced host translation shutoff recovers approximately 20-30% of the cellular energy that is expended in translating host mRNAs (Buttgereit and Brand 1995), subverting this energy for viral protein synthesis. Several viruses, including vaccinia virus, utilize this host shutoff pathway to enhance viral protein synthesis(Abernathy and Glaunsinger 2015).

Cells combat viral infection through a variety of host defense responses aimed at limiting viral protein production by triggering innate immune signaling pathways(Jan et al. 2016; Stern-Ginossar et al. 2019). For example, one cellular response to proteotoxic stress, including viral infection, is the activation of the integrated stress response (ISR) resulting in eIF2α phosphorylation and the subsequent global repression of translation initiation(Costa-Mattioli and Walter 2020; Liu et al. 2020).

Cells utilize the eIF2α kinase, double-stranded RNA (dsRNA)-dependent protein kinase (PKR), to inhibit protein biogenesis in the presence of viral RNA(McCormack et al. 1992; Williams 1999). As viral propagation requires the cellular translation machinery, viruses have evolved a myriad set of strategies to combat such host defense mechanisms(Jan et al. 2016). For example, vaccinia virus, a large double stranded DNA virus that replicates entirely in the cytoplasm of the infected cells(Moss 2013), encodes two different proteins that either bind and occlude double-stranded RNA detection by PKR, or act as a PKR pseudo-substrate(Chang et al. 1992; Davies et al. 1992; Carroll et al. 1993; White and Jacobs 2012; Meade et al. 2019). The combined impact of these viral proteins is to limit host translation shutdown, highlighting the requirement for sustained ribosome output for rapid viral protein production. These observations also suggest that viral replication requires sustained high levels of active ribosomes to mediate rapid translation of viral mRNAs.

Actively translating ribosomes can become stalled for a sufficient length of time to allow for a trailing ribosome to collide with the stalled ribosome. These ribosome collisions trigger the ribosome-associated quality control (RQC) pathway that acts to ubiquitylate and destroy the ribosome-associated nascent chain as well as catalyze ribosome subunit splitting and mRNA disengagement (Joazeiro 2019; Inada 2020). These outcomes limit the abundance of potential toxic truncated translation products and recycle ribosomes to reenter the translation cycle. The inability to recognize collided ribosomes and initiate the RQC pathway results in ribosomal readthrough of stall inducing sequences and codon frameshifting(Juszkiewicz and Hegde 2017; Sundaramoorthy et al. 2017; Wang et al. 2018; Joazeiro 2019; Juszkiewicz et al. 2020a). Additionally, a failure to recycle collided ribosomes can deplete the active ribosome pool and result in reduced tissue function(Mills et al. 2016; Mills and Green 2017; Liakath-Ali et al. 2018). These observations suggest that cellular conditions that demand constant protein production in the face of enhanced ribosome collisions, such as those encountered upon viral infection, may require RQC function to maintain sufficient levels of available ribosomes.

In mammals, the RQC pathway requires site-specific regulatory ribosomal ubiquitylation (RRub), catalyzed by the ubiquitin ligase ZNF598 (Juszkiewicz and Hegde 2017; Matsuo et al. 2017; Sundaramoorthy et al. 2017). ZNF598 ubiquitylates the 40S ribosomal proteins eS10 and uS10 at precise lysine residues and either loss of ZNF598 or mutations that block eS10 or uS10 ubiquitylation result in RQC failure and ribosomal readthrough of stall-inducing sequences(Juszkiewicz and Hegde 2017; Sundaramoorthy et al. 2017; Garshott et al. 2020). Thorough biochemical and genetic studies in multiple organisms have delineated a growing list of molecular constituents within the RQC pathway that act on collided ribosomes(Joazeiro 2019; Inada 2020). Recent studies have uncovered cellular signaling pathways that are induced by collisions that both mount a transcriptional response and result in translation initiation inhibition by multiple routes(Tollenaere et al. 2019; Hickey et al. 2020; Juszkiewicz et al. 2020a; Meydan and Guydosh 2020; Sinha et al. 2020; Vind et al. 2020; Wu et al. 2020). How viruses utilize various RQC components for viral propagation or are impacted by RQC dysfunction remains largely unexplored.

Here we utilize vaccinia virus, which encodes ~200 viral proteins, to interrogate the interplay between the RQC pathway and viral protein synthesis. We demonstrate that ZNF598 and uS10 ubiquitylation is needed for optimal vaccinia virus replication. Chronic loss of ZNF598 function resulted in wide-spread reduction of nearly all viral protein production with little effect on viral transcription. Further, vaccinia virus infection reduced overall cellular RQC activity, indicating that viral protein production depletes a critical RQC factor. Taken together, we conclude that vaccinia virus infection induces ribosome collisions and that the RQC pathway, through ZNF598 and its substrate uS10, rescues stalled ribosomes from non-functional translation events and recycles ribosomal subunits for optimal translation of viral mRNAs.

## Results

### Ribosome collisions occur during integrated stress response activation in the absence of eIF2α phosphorylation

Viral replication consumes the host translation machinery and host defense strategies often target the translation machinery to combat viral propagation(Jan et al. 2016). The observation that viruses often employ countermeasures to maintain high rates of translation suggest that any loss in available ribosomes may be particularly deleterious to viral replication. RQC pathway activation upon ribosome collisions acts to recycle stalled ribosomal subunits for reuse during translation(Joazeiro 2019). As such, viruses may depend on the RQC machinery to liberate collided and stuck ribosomes during rapid translation of viral mRNAs. Many viruses, including vaccinia virus, antagonize PKR-dependent eIF2α phosphorylation to prevent host shutdown of translation initiation(Meade et al. 2019; Dhungel et al. 2020). To examine if ribosome collisions occur at a higher frequency during innate immune or ISR activation when eIF2α phosphorylation is compromised, we utilized mouse embryonic fibroblasts (MEFs) containing either wild type or S51A mutant eIF2α that cannot be phosphorylated(Scheuner et al. 2001). We used polyI:C transfection to mimic viral infection mediated activation of innate immune signaling pathways. PolyI:C transfection resulted in robust eIF2α phosphorylation in WT MEFs and failed to induce eS10 or uS10 ubiquitylation which are markers for ribosome collisions (Fig. 1A). In contrast, polyI:C transfection in cells unable to repress translation through eIF2α phosphorylation resulted in a time-dependent increase in uS10 ubiquitylation indicating an induction of ribosome collisions (Fig. 1A). To further examine the impact of compromised eIF2α phosphorylation on ribosome collisions we exposed cells to UV which induces ribosome collisions and ISR. As expected, UV exposure resulted in ZNF598-dependent eS10 and uS10 ubiquitylation despite enhanced eIF2α phosphorylation (Garshott et al. 2020). However, eS10 and uS10 ubiquitylation was enhanced in eIF2a mutant MEFs further indicating that an inability to repress translation initiation upon proteotoxic stress increases ribosome collision frequency (Fig. 1B). In human cells, addition of ISRIB, which blocks the inhibitory activity of phospho-eIF2α (Sidrauski et al. 2015), enhanced the UV-dependent eS10 and uS10 ubiquitylation (Fig. 1C) and indicates that an inability to downregulate translation via eIF2α phosphorylation results in an elevated frequency of ribosome collisions upon UV exposure. Taken together, these results suggest that antagonizing eIF2α phosphorylation during proteotoxic stress enhances ribosome collisions.

**Figure 1.**
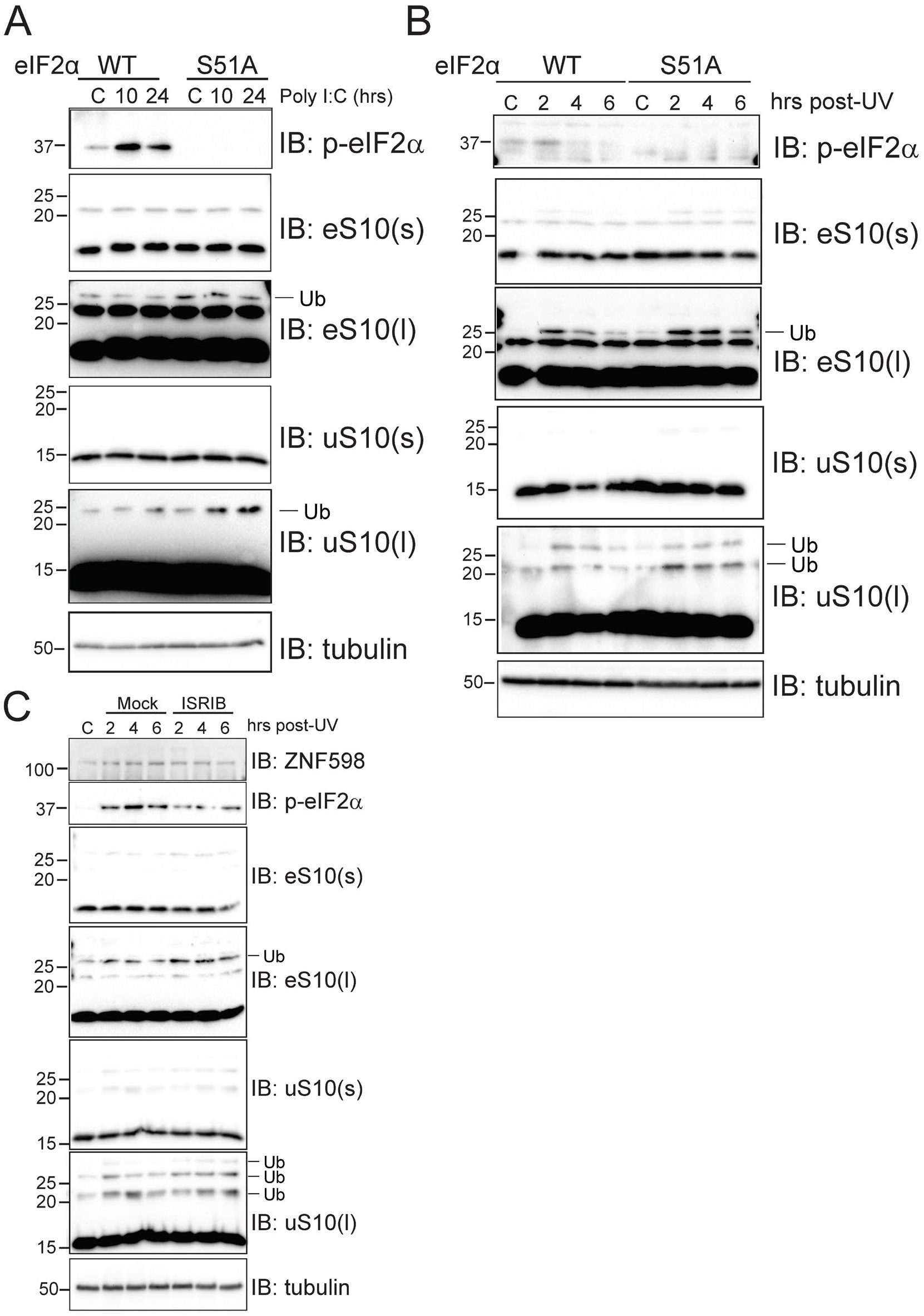
Loss of eIF2α phosphorylation leads to higher frequency ribosome collisions during proteotoxic stress. A) Mouse embryonic fibroblasts (MEFs) derived from wild type (WT) or S51A mutant eIF2α animals were mock transfected (C) or transfected with polyI:C and cells were harvested at the indicated time points post transfection. Whole cell extracts were analyzed by SDS-PAGE and immunoblotted with the indicated antibodies. (S) and (L) denote short and long exposures, respectively. B) MEFs of the indicated genotypes were unexposed (C) or exposed to 200J/m^2^ UV and allowed to recover for the indicated time prior to harvesting. Cells were harvested at the indicated time-points, whole cell extracts were run on SDS-PAGE and immunoblotted with the indicated antibodies. C) 293T cells were exposed to 200J/m^2^ UV and allowed to recover in normal media or in the presence of 200nM ISRIB. Cells were harvested at the indicated timepoints and total lysates were resolved on SDS-PAGE and immunoblotted using the indicated antibodies.

### Vaccinia virus replication is inhibited upon loss of ZNF598 or uS10 ubiquitylation

We reasoned that increased the proteotoxic stress from viral infection, combined with viral antagonism of eIF2a phosphorylation, might make RQC particularly important for viruses. To directly test if vaccinia virus replication is impacted by loss of RQC activity, parental and ZNF598 knockout (KO) cells were infected with vaccinia virus and viral titers were determined. vaccinia virus replication was repressed more than 10-fold in two independent ZNF598 KO cell types which is consistent with previous results(DiGiuseppe et al. 2018) (Fig. 2A). We previously generated and characterized point mutant knock-in cell lines in which critical lysine residues (K138,K139 within eS10 and K4,K8 within uS10) in eS10 or uS10 that are ubiquitylated by ZNF598 and are required for RQC activity were mutated to arginine (Garshott et al. 2020). Vaccinia virus infections in mutant eS10 or uS10 cell lines resulted in 10-100 fold less viral replication compared to parental cells (Fig. 2B). Interestingly, multiple clones of uS10 point mutant lines displayed a more profound defect in vaccinia virus replication compared to the eS10 mutant cell line across several experiments. These results confirm that vaccinia virus requires ZNF598-mediated ribosomal ubiquitylation for optimal viral replication(DiGiuseppe et al. 2018). Consistent with the viral titer data, vaccinia virus infection resulted in enhanced uS10 ubiquitylation throughout the infection time course indicating that vaccinia virus infection induces ribosome collisions during cellular replication (Fig. 2C). While eS10 ubiquitylation was also stimulated at early time points, it was less consistently ubiquitylated compared to uS10 upon vaccinia virus infection. Previous studies demonstrated that ZNF598 utilizes an internal poly proline motif to engage in repression of AU rich cytokine transcripts by binding to the GIGYF-4EHP complex (Tollenaere et al. 2019). ZNF598 also represses expression of interferon stimulated genes (ISGs), which are critical for pathogen defense responses (DiGiuseppe et al. 2018). It is noteworthy that these ZNF598 functions do not depend on its ligase activity. Transient loss of ZNF598 was shown to be broadly antiviral due to the derepression of anti-inflammatory cytokines(DiGiuseppe et al. 2018; Tollenaere et al. 2019). We monitored ISG stimulation upon IFN-β addition to examine if interferon signaling was constitutively activated in our RQC mutant cell lines. The basal levels of the known ISG ISG15 were not elevated in ZNF598 KO cells or in our ribosome point mutation knock in cells compared to parental cells (Fig. 2D,E). Moreover, ISG expression in these RQC deficient cells lines mirrored WT cells, with IFN-β addition resulting in robust ISG15 production in all cell lines (Fig. 2D,E). These results, combined with the observation that vaccinia virus replication is mostly insensitive to interferon addition(Smith et al. 2018), argue that the loss of RQC pathway function is responsible for the observed vaccinia virus replication defect in ZNF598 KO cell lines.

**Figure 2.**
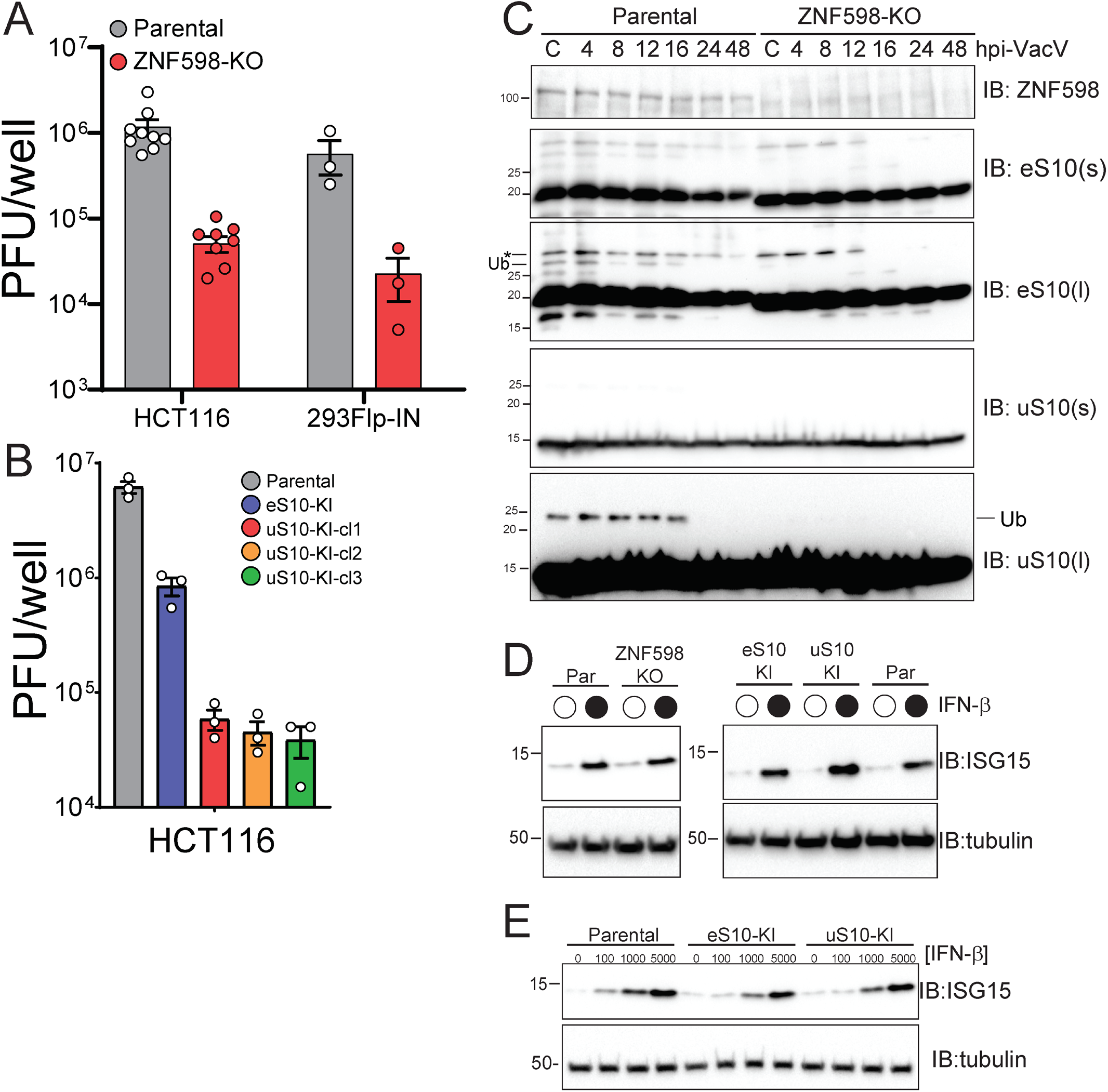
Vaccinia virus replication is attenuated in cells unable to mount an RQC response. A) ZNF598 knockout (KO) or parental HCT116 or 293Flp-IN cells were infected with vaccinia virus (VacV) (10,000 PFU/well (MOI=~0.04)). Cells were collected 24 hours post infection and viral titers were determined by plaque assay. B) Cells containing mutant ribosomal proteins eS10 (K138R;K139R-eS10-KI), uS10 (K4R;K8R-uS10-KI), or parental cells were infected with vaccinia virus (10,000 PFU/well (MOI=~0.04)). Cells were collected 24 hours post infection and viral titers were determined by plaque assay. Three individual clones (cl) of uS10-KI cells were tested. C) Parental or ZNF598 knockout (KO) HCT116 cells were mock infected (C) or infected with VacV (MOI=5) and cells were collected at the indicated time points and analyzed by SDS-PAGE and immunoblotted with indicated antibodies. D,E) The indicated cell lines were either untreated or treated with 1000U of IFN-β for 16 hours (D) or treated with the indicated concentration of IFN-β for 16 hours and cells were harvested and analyzed by SDS-PAGE.

### ZNF598 mediated uS10 ubiquitylation is needed for optimal vaccinia virus protein synthesis

Based on our data and previous reports, it is possible that translation of a specific subset of essential vaccinia virus mRNAs are specifically inhibited upon loss of ZNF598-mediated uS10 ubiquitylation(DiGiuseppe et al. 2018). Alternatively, defective RQC pathway activity could broadly impact ribosome availability which would result in reduced translation of many viral mRNAs. To differentiate between these possibilities, we performed stable isotope labelling of amino acids in culture (SILAC) based quantitative proteomics experiments upon infection with vaccinia virus in parental and ZNF598 knockout cells. Heavy (Lys8) labeled parental or ZNF598 knockout cells were infected with vaccinia virus at an MOI=5 and cells were harvested at seven timepoints post infection and mixed with light labeled uninfected cells of the same genotype (Fig. 3A). We quantified the abundance of 64 viral proteins in at least a single time point (Table S1). As expected, viral protein abundance increased over time with the most viral proteins detected 48 hours after infection (Fig. 3B,C). An observable reduction in the mean vaccinia virus protein abundance in ZNF598 KO cells compared to parental controls could be detected 4 hours post infection and reached statistical significance at all time points after 16 hours (Fig. 3C,D). No difference in overall host protein abundance was observed between ZNF598 KO and parental cells (Fig 3D, Table S1). These results are consistent with our viral titer assays in that viral protein production is repressed in ZNF598 KO cells. To verify that this result was due to an inability to mount an RQC response, we performed a similar proteomics experiment in parental and uS10 point mutant knock in cells (uS10-KI). We modified the labeling scheme for this series of experiments in that heavy labeled uS10-KI cells and light parental cells were both infected at an MOI=5 and the two cellular genotypes were collected over a similar time course post infection and mixed (Fig. 3A). As observed in ZNF598 KO cells, overall host protein abundance was unchanged upon infection, but vaccinia virus protein production was broadly repressed in uS10-KI cells (Fig. 3E-G). Consistent the previous reports, a small number of host proteins displayed vaccinia virus and genotype specific abundance changes (Soday et al. 2019). However, despite these changes, overall host protein abundance was not impacted by vaccinia virus infection (Fig. 3G). Combined, our data suggests that an inability to ubiquitylate collided ribosomes results in a functional ribosomal imbalance, leading to compromised viral protein production.

**Figure 3.**
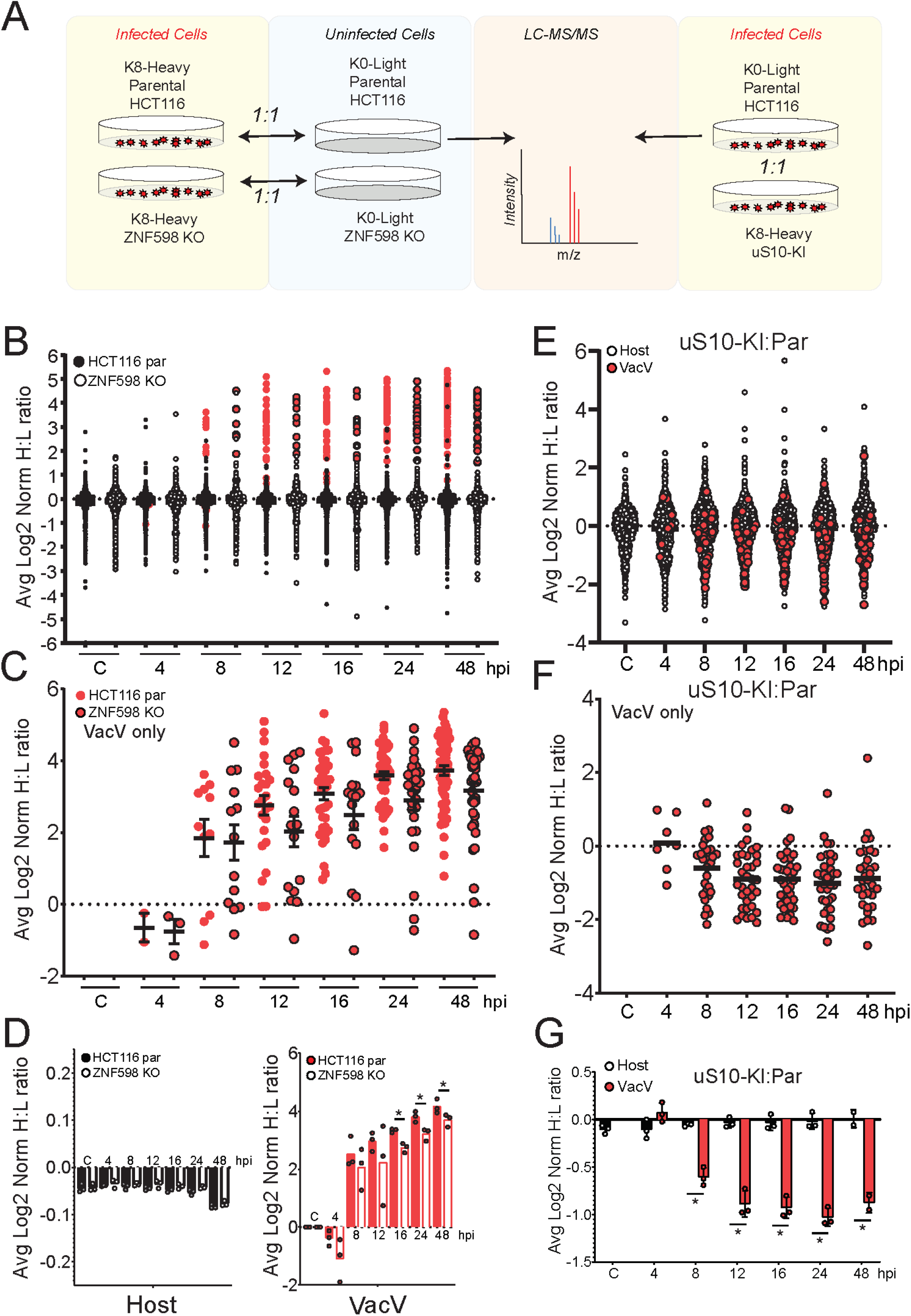
Proteomic characterization of vaccinia virus protein production in RQC deficient cells. A) Schematic showing the labelling strategy for the SILAC experiments. B) Heavy-labeled parental (filled circles) or ZNF598 KO cells (open circles) were infected with vaccinia virus (MOI=5), collected at the indicated time points post infection, and mixed 1:1 with light-labeled uninfected cells of the same genotype. Samples were analyzed by mass spectrometry and normalized log2 heavy to light ratios were determined for host (black) and vaccinia virus (VacV) (red) proteins. The average normalized log2 H:L ratio for proteins across three replicate experiments is depicted. C) Normalized log2 heavy to light ratios of VacV proteins from parental (red circles, no border) or ZNF598 KO (red circles, black border) cells infected with VacV (MOI=5) at the indicated time points. The average normalized log2 H:L ratio for proteins across three replicate experiments is depicted. Black bars denote the mean of all VacV H:L ratios at a given time point with error bars representing SEM. D) The average normalized log2 H:L ratio for host (left) or VacV (red, right) proteins in parental (filled black or red bars) or ZNF598 KO (unfilled bars) cells at each time point. Error bars denote SEM for triplicate experiments. *-pval<0.05 – one-way ANOVA. E,F) Light-labeled parental (closed circles) or heavy-labeled uS10-knock in (KI) cells (open circles) were infected with vaccinia virus, collected at the indicated time points post infection, and mixed 1:1. Samples were analyzed by mass spectrometry and normalized log2 heavy to light ratios were determined for host (black) and VacV (red) proteins (E) or just VacV proteins (F). The average normalized log2 H:L ratio for proteins across three replicate experiments is depicted. Black bars denote the mean of all VacV H:L ratios at a given time point. G) The average normalized log2 H:L (parental:uS10-KI) ratio for host (black bars) or VacV (red bars) proteins at each time point. Error bars denote SEM for triplicate experiments. *-pval<0.05 – one-way ANOVA.

### Translation of mRNAs containing a 5’UTR polyA sequence is not specifically impacted by loss of ZNF598 activity

Consistent with previous reports, we demonstrate that vaccinia virus replication is reduced in cells that lack an ability to ubiquitylate ribosomes during RQC events(DiGiuseppe et al. 2018). However, contrary to previous reports suggesting that translation of late vaccinia virus mRNAs containing non-templated polyA sequences of various lengths within their 5’UTR were specifically impacted by loss of ZNF598 activity, we demonstrate that viral protein production is broadly repressed in ZNF598 KO or uS10-KI cells. Direct comparison of viral protein abundance with and without reported polyA sequences within the 5’UTR of corresponding mRNAs(DiGiuseppe et al. 2018) did not reveal any difference in either ZNF598 KO or uS10-KI cells (Fig. 4A,B). This result suggests that viral protein production is repressed irrespective of the presence of 5’UTR polyA sequences in RQC deficient cells. To probe this hypothesis further, we constructed reporter plasmids containing polyA sequences of varying lengths immediately prior to the start codon of the firefly luciferase coding sequence. Consistent with previous reports, firefly luciferase protein production decreased as the 5’UTR polyA sequence length increased as compared to a control renilla luciferase reporter (Fig. 4C) (Dhungel et al. 2017). Performing a similar experiment in vaccinia virus infected cells revealed the same polyA length dependent decrease in firefly luciferase protein production. However, vaccinia virus infection resulted in enhanced firefly luciferase protein levels with the longest polyA sequence compared to uninfected cells (Fig. 4C). This is consistent with previous reports showing that viral mediated phosphorylation of the 40S ribosomal protein RACK1 facilitates translation of late viral mRNAs with 5’UTR polyA sequences(Jha et al. 2017). Transfection of these reporters into ZNF598 KO, uS10-KI, or eS10-KI cells resulted in a similar polyA-dependent repression of firefly luciferase protein levels indicating that a loss of RQC activity does not allow for enhanced protein production for mRNAs containing a 5’UTR polyA sequence (Fig. 4D). Previous studies demonstrated that loss of ZNF598 results in enhanced protein production downstream of polyA sequences within the coding sequence of reporter mRNAs(Juszkiewicz and Hegde 2017; Sundaramoorthy et al. 2017). This result is due to an inability to sense collided ribosomes during translation elongation. RQC pathway components have not been demonstrated to play a role during translation initiation and our data indicates that mRNAs containing non-coding polyA sequences within the 5’UTR, while broadly repressive, are not translated more efficiently in cells with defective RQC activity.

**Figure 4.**
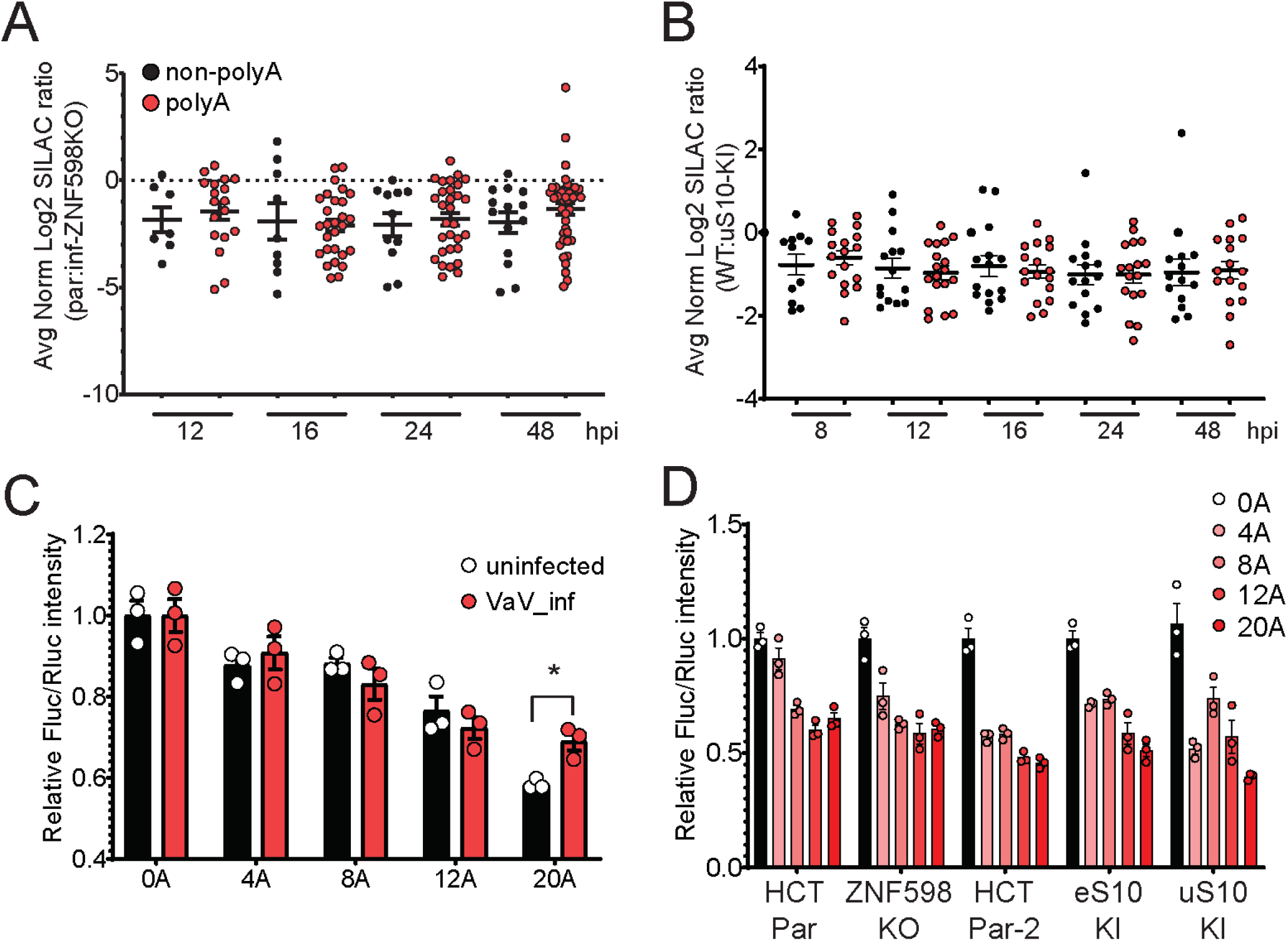
RQC deficiency does not specifically impact translation of mRNAs containing a 5’UTR polyA sequence. A) The difference in the normalized Log2 H:L ratios for vaccinia virus (VacV) proteins with (red circles) or without (black circles) annotated polyA sequences within the 5’UTR of corresponding mRNAs between infected parental and ZNF598-KO cells is depicted. Black bars denote the mean of all H:L ratios at a given time point with error bars representing SEM. B) The normalized Log2 H:L (parental:uS10-KI) ratios of VacV proteins with (red circles) or without (black circles) annotated polyA sequences within the 5’UTR of corresponding mRNAs. Black bars denote the mean of all H:L ratios at a given time point with error bars representing SEM. C) Relative luminescence intensities of firefly luciferase reporters containing the indicated length of polyA sequence immediately before the start codon for firefly luciferase compared to renilla luciferase controls without polyA sequences is shown for uninfected (black bars) and VacV infected (red bars) cells. Error bars denote SEM for triplicate experiments. *-pval<0.05 – Student’s t-test. D) Relative luminescence intensities of firefly luciferase reporters containing the indicated length of polyA sequence as compared to renilla luciferase controls is shown for transfections into the indicated cell types. Error bars denote SEM for triplicate experiments.

### Dual-RNA-seq of vaccinia infection reveals host and vaccinia transcriptional response to be RQC independent

It is possible that our proteomics results demonstrating broad repression of viral protein production in RQC deficient cell lines is due to defects in viral mRNA transcription. To directly address this possibility, we performed RNA-seq analysis of mock treated and vaccinia virus infected parental or uS10-KI cells at 4- and 8-hours post-infection (Fig. 5A). The sequencing experiment generated approximately 15 million reads per sample per timepoint, totaling 270 million reads (Fig. S1A). Approximately 12,000 human genes were quantitatively documented across each timepoint and 200 vaccinia genes were sampled in infected samples (Table S2). The reads were aligned in parallel to human and Vaccina genomes to investigate the transcriptional dynamics during infection. Mock treated samples aligned >97% reads to the human genome while vaccinia virus infected cells, 4 hours post infection, aligned 91-93% of reads to the host genome (Fig. 5B). In parental cells. the percentage of reads mapped to the host genome further decreased to 59% at 8 hours post infection demonstrating the expected increase in viral mRNA abundance during infection (Fig. 5B). The reads aligned to vaccinia genome increased from 6% at 4hrs to 39% 8 hours post infection in parental cells and from 4% to 26% in uS10-KI cells (Fig. 5C). Prior reports in Hela cells infected with vaccinia virus identified 70% of mRNA at 8 hours post infection to be of viral origin(Yang et al. 2015).

**Figure 5.**
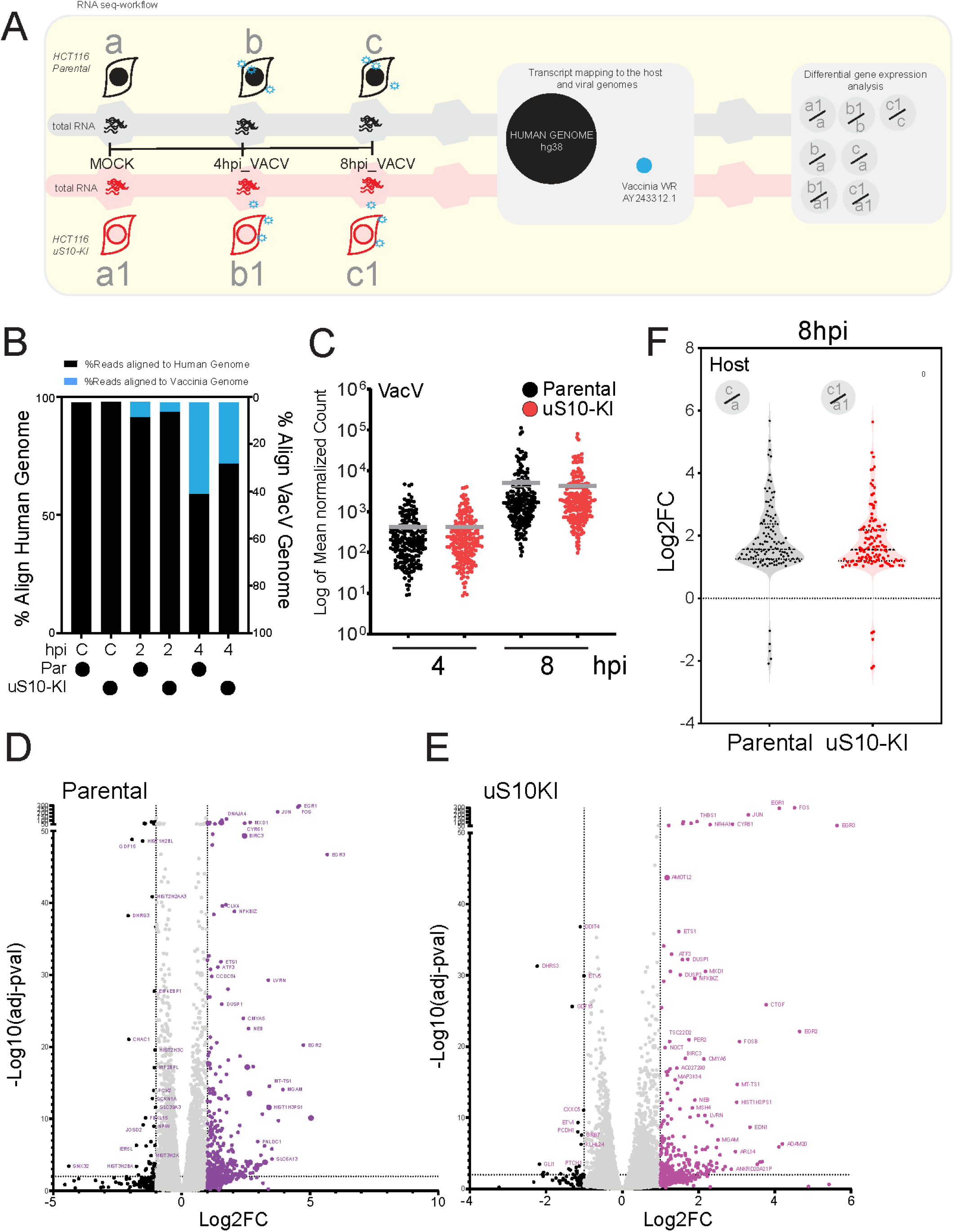
RNA-seq identifies intact viral and host transcriptional response in uS10-KI cells. A) A schematic depicting the RNA-seq experimental design and analysis. Individual RNA library preparation time-points are depicted in (a,b,c) or (a1,b1,c1) for Parental and uS10-KI cells, respectively. B) Percent mRNA alignment to the human (black bars) or vaccinia virus (VacV) (blue bars) genome in parental or uS10-KI cells. C) Reads aligned to the VacV genome were analyzed by DESeq2 and the log transformed mean normalized counts of 200 VacV genes is depicted for infected parental (black dots) or uS10-KI (red dots) cells at 4 and 8 hours post infection. Grey bars indicate mean and error bars indicate SEM across triplicate samples. D,E) Volcano plot showing differentially expressed host genes 8 hours post infection in parental (D) or uS10-KI (E) cells. Colored dots indicate statistically significant induced genes (relative to uninfected cells), while black dots indicated repressed genes. F) Violin plot showing frequency distribution of the log2 ratios of the 122 common differentially expressed host genes comparing uninfected cells to those 8 hours post infection (hpi) within the indicated cell lines.

We analyzed the transcriptional dynamics of vaccinia genes at 4 and 8 hours post infection within parental or uS10-KI cells. We observed no difference in vaccinia virus mRNA levels 4 hours post infection between parental and uS10-KI cells (Fig. 5C). There was a small, but noticeable overall decrease in the mean vaccinia virus mRNA abundance in uS10-KI cells compared to parental cells 8 hours after vaccinia virus infection (Fig. 5C). This decrease is consistent with our data showing a defect in vaccinia virus viral propagation in uS10-KI cells. Analysis of the differentially expressed genes between parental and uS10-KI cells revealed 39 vaccinia virus mRNAs (out of 200) with reduced expression in uS10-KI cells (Table S2). Overall, our analysis indicates that while a small number of vaccinia virus genes show reduced mRNA levels at 8 hours post infection, there is no evidence of widespread alternations in vaccinia virus gene transcription in uS10-KI cells (Figure S1B, Table S2). These results are consistent with a model in which the observed difference in viral protein production in our proteomics experiments is due to a post-transcriptional mechanism.

We next analyzed the host transcriptional response to vaccinia infection and compared gene expression changes as a function of viral infection within the same cell type. Clear host transcriptional responses can be observed 8 hours after infection including the documented MAPK-based activation of FOS, EGR1, and c-JUN expression in both parental and uS10-KI cells (Fig. 5D,E). At 8 hours post infection, we observed 332 and 228 host genes with differential expression compared to uninfected controls in parental and uS10-KI cells, respectively (Table S3). Interrogation of the 122 genes that overlapped between these two gene sets revealed a similar distribution of differential expression between the two cell types (Fig. 5F, S1C-E). Overall, the host transcriptional response to vaccinia virus infection is largely unaltered in uS10-KI cells compared to parental controls. Combined, the RQC deficient uS10-KI cells displayed similar host and viral transcript levels upon vaccinia virus infection. Given the wide fluctuation in viral proteins, we propose that a post-transcriptional mechanism governs the loss of viral protein expression.

### Vaccinia virus infection limits cellular RQC activity

Combined, our data suggests that vaccinia virus infection enhances the frequency of ribosome collisions and an inability to clear these collisions compromises vaccinia virus mRNA translation. These results support a model in which vaccinia virus replication relies on a cellular pool of available ribosomes that becomes depleted in RQC defective cells. To probe this possible model, we utilized a well-characterized fluorescent RQC reporter in which the genes encoding GFP and ChFP are separated by a third coding sequence containing a decoded polyA sequence (Fig. 6A)(Juszkiewicz and Hegde 2017; Sundaramoorthy et al. 2017). Cells with reduced ZNF598 protein expression or containing eS10 or uS10 mutations in ubiquitin-acceptor lysine residues display enhanced ChFP levels downstream of the polyA sequence relative to upstream GFP levels(Juszkiewicz and Hegde 2017; Sundaramoorthy et al. 2017; Garshott et al. 2020). vaccinia virus infection of cells expressing a control reporter lacking a polyA sequence resulted in a time dependent decrease in both GFP and ChFP expression indicative of host dependent translation repression upon vaccinia virus infection (Fig 6B). Infection of cells expressing the polyA-containing stall reporter also resulted in translational repression of GFP expression. However, vaccinia virus infection also resulted elevated ChFP expression relative to GFP 16 hours post infection (Fig. 6B). This result suggests that cellular RQC activity is decreased upon vaccinia virus infection and that vaccinia virus viral factories may be titrating ZNF598 activity away from the cellular pool. Taken together, our data are consistent with a model in which vaccinia virus viral protein production results in ribosome collisions that must be cleared to recycle ribosomes back into the available pool to maintain high vaccinia virus mRNA translation activity required for optimal viral production.

**Figure 6.**
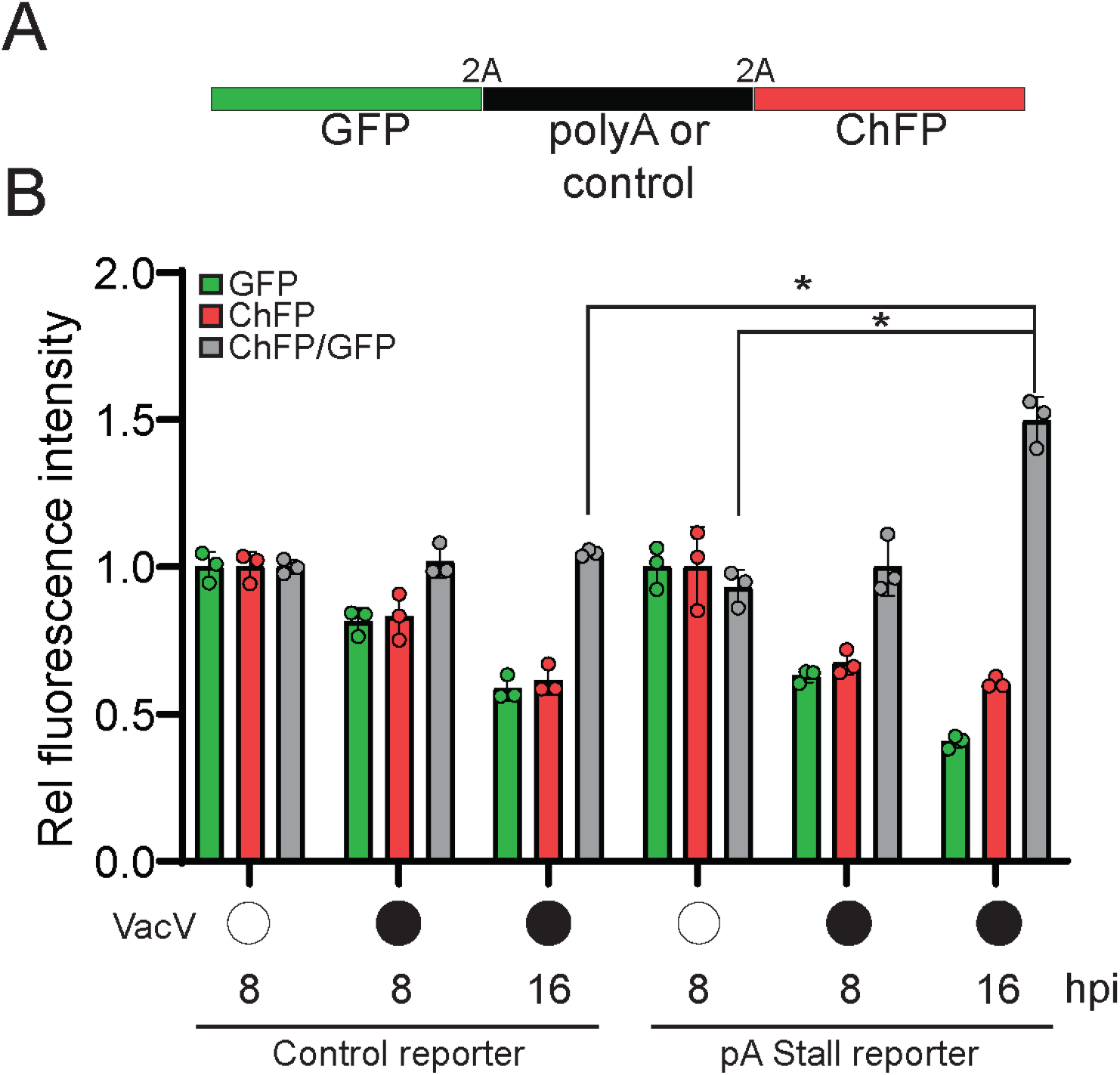
Vaccinia virus infection alters cellular RQC activity. A) Schematic of fluorescent stall reporter plasmid with either a control or polyA sequence in the middle coding sequence. 2A denotes ribosomal skipping sequence. B) The GFP, ChFP, or ChFP/GFP ratio in cells transfected with the either control or polyA-containing stall reporter plasmids is depicted for uninfected or vaccinia virus (VacV) infected cells at the indicated time points post infection. Error bars denote SEM for triplicate experiments. *-pval<0.05 – Student’s t-test.

## Discussion

Viruses are entirely dependent on the host for translation of their proteins. Indeed, several mechanisms allow viruses to efficiently hijack cellular machinery for optimal viral translation initiation, elongation and termination(Jan et al. 2016). However, several different host mechanisms have also evolved to limit viral protein translation, including PKR-mediated eIF2α phosphorylation as a means to prevent viral mRNA translation initiation(Stern-Ginossar et al. 2019). These viral and host adaptations reveal the importance of translation control for the ongoing conflict between viruses and the hosts they infect.

Less well studied is the role that RQC factors may play in viral infection and host control of viral infections. Recent discoveries have revealed the important regulatory role RQC plays during host translation especially under conditions the elevate the frequency of ribosome collisions(Joazeiro 2019). A large number of evolutionarily conserved proteins coordinate RQC within a signaling hierarchy(Sitron and Brandman 2020). The physiological role for RQC factors in maintaining ribosome availability is exemplified by Pelota, which assists in recycling stalled ribosomal subunits(Pisareva et al. 2011; Guydosh and Green 2014), and where loss-of-function results in diminished translation of globin mRNAs in blood cells (Mills et al. 2016). Despite these studies, the physiological relevance of many RQC components remains uncharacterized. Of particular interest is the E3 ubiquitin ligase ZNF598, which ubiquitylates 40S ribosomal proteins and recruits downstream ribosome rescue factors such as the RNA helicase ASCC3(Matsuo et al. 2017; Hashimoto et al. 2020; Juszkiewicz et al. 2020b). The important role that RQC plays in resolving collided ribosomes and other stalled ribosomes during proteotoxic stress suggests that these factors may become particularly important during viral infection. Indeed, while ribosomes are among the most abundant cellular machines in nearly all cells, RQC factors are several fold less abundant than ribosomes(Itzhak et al. 2016), suggesting that a large burden on the RQC system could possibly overwhelm its capacity.

To address the physiological importance of RQC activity, we investigated its role during viral infection and the host response. We focused on vaccinia virus, as this virus is known to substantially redistribute ribosomes for selective translation of viral mRNAs as well as prevent PKR-mediated eIF2α phosphorylation. Consistent with increased translation stress, our work shows RQC is activated during vaccinia virus infection. Similar to virus-induced proteotoxic stress, we observed increased translation stress upon exposure to polyI:C or UV that is revealed by disabling eIF2α phosphorylation. These data led us to test whether RQC may become limiting during vaccinia virus infection by disrupting ZNF598 or one of its primary ubiquitylation targets on the 40S ribosomal subunit, uS10. Consistent with previous work(DiGiuseppe et al. 2018), we found that ZNF598 and its uS10 target are required for optimal vaccinia virus replication. Although previous work has indicated that ZNF598 is important for translation of polyA-containing vaccinia proteins(DiGiuseppe et al. 2018) and can serve as a negative regulator of the interferon response(Tollenaere et al. 2019), we find that neither increased interferon activation nor selective loss of polyA-containing proteins can explain the decrease in viral replication. Instead, by performing quantitative proteomic and RNAseq analyses after vaccinia virus infection, we find that there is a strong post-transcriptional blockade to protein synthesis of nearly every viral protein while leaving the host proteome relatively unchanged. These results are thus consistent with an increased reliance on RQC pathways during vaccinia virus infection to clear collided ribosomes that result from the dual insults of increased proteotoxic stress and antagonism of eIF2α-mediated translation arrest.

Based on our data we propose the following model for how RQC is impacted and plays a functional role during vaccinia virus infection (Fig 7). Vaccinia virus infection results in both an increased burden on the translation machinery, as well as a suppression of the cellular integrated stress response (ISR) through antagonism of eIF2α phosphorylation. Both of these consequences of vaccinia virus infection aggravate ribosome collisions. In wild type cells, RQC activation upon ribosome collisions rescues collided ribosomes, allowing re-entry of these ribosomes into the pool of ribosomes available for active translation. This replenished pool of ribosomes allows optimal viral protein synthesis. However, in RQC deficient cells, either through loss of ZNF598 or mutation of the primary ubiquitylation target on uS10, vaccinia virus induced ISR suppression and ribosome collisions are no longer resolved by the RQC pathway. As a result, a cycle of dysfunctional ribosome collisions leads to a depleted ribosome pool and a decrease in synthesis of nearly every viral protein.

**Figure 7.**
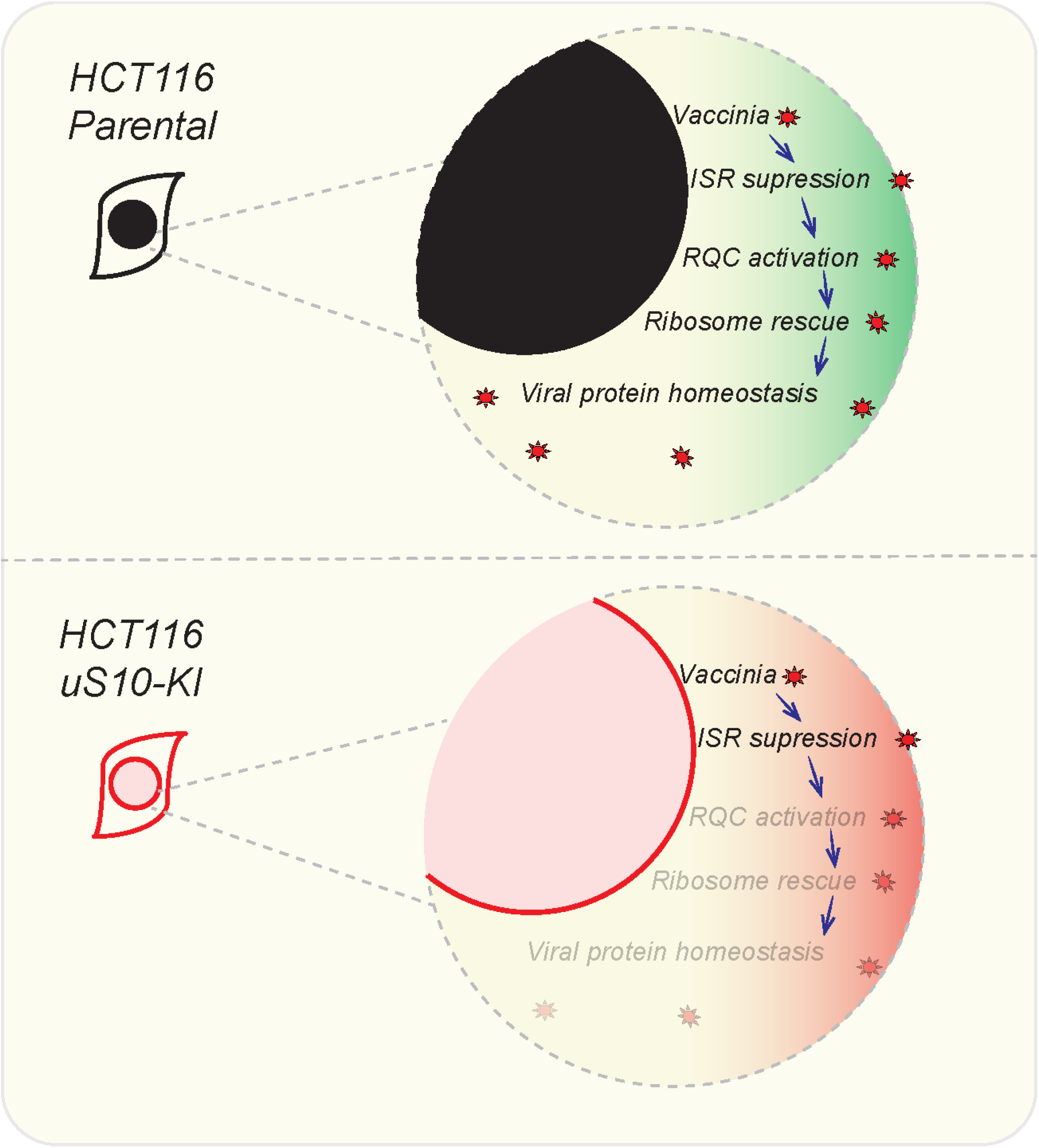
Model of RQC activation and ribosome depletion during vaccinia virus replication: In the upper panel, vaccinia virus infection in parental cells and subsequent ISR suppression results in RQC activation and ribosome rescue with normal viral protein synthesis. In the lower panel, ISR suppression by vaccinia virus in RQC defective cells, induces futile ribosome collisions and loss of ribosome rescue.

Our work indicates that the evolution of a viral response to the host ISR has exposed the virus to a conditional requirement for the RQC pathway in regulating translational reprogramming. While ribosome collisions are part of the regular translation cycle, feedback inhibition of translation initiation through the ISR normally curtails overt RQC activation(Hickey et al. 2020; Juszkiewicz et al. 2020a; Sinha et al. 2020; Vind et al. 2020; Wu et al. 2020). When viruses interfere with this feedback inhibition, such as vaccinia virus antagonism of PKR-mediated eIF2α phosphorylation, non-productive translation cycles ensue and therefore require RQC based ribosomal rescue. We suspect several different viruses that alter ISR response might be conditionally dependent on the RQC pathway for propagation.

## Acknowledgements

This work was supported by funding from the National Institutes of Health (R35 GM133633, M.D.D.), (DP2-GM119132, E.J.B.), (T32 GM007240, A.P.R.), Pew Biomedical Scholars Program (M.D.D.), and Hellman Fellows Program (M.D.D.).

## Materials and Methods

### Cell lines

HCT116 (ATCC), HEK293T (ATCC) and MEFs (WT and S51A) cells (Scheuner et al. 2001) were grown in DMEM ((high glucose, pyruvate and L-Glutamine) media supplemented with 10% FBS and maintained in humidified incubator supplied with 5% CO2 atmosphere. ZNF598 knockout cells were validated and characterized previously (Sundaramoorthy et al. 2017). uS10 knock-in (uS10-KI(K4R/K8R) and eS10 knock-in (eS10-KI(K138RK139R) HCT116 cells were generated and characterized previously (Garshott et al. 2020).

### Reagents and Antibodies

The following chemical reagents were used at the indicated concentrations: ISRIB (Tocris) 200nM, DTT (Sigma-Aldrich) 5mM. Cells were irradiated using Spectrolink UV-crosslinker using energy setting (200J/m2). The following antibodies were used rabbit polyclonal RPS10/eS10 (Abclonal-A6056), RPS2/uS5 (Bethyl-A303-794A), RPS3/uS3 (Bethyl-A303-840A), Rabbit polyclonal ISG15 (Cell Signaling Technology-2743)Rabbit polyclonal anti-ZNF598 (Sigma-Aldrich-HPA041760) rabbit monoclonal RPS20/uS10 (Abcam-ab133776), Phospho-eIF2α (Ser51) (Cell Signaling Technology-D9G8) Mouse monoclonals used : α-Tubulin (Cell Signaling Technology DM1A). The following HRP conjugated secondary antibodies were used Anti-Rabbit IgG (H+L), HRP Conjugate antibody (Promega -W4011), Anti-Mouse IgG (H+L), HRP Conjugate Antibody (Promega-W4021). Immuno-blots were developed using enhanced chemiluminescence (Clarity -Biorad) on a XRS+ biorad imager. Cells were transfected using Lipofectamine RNAiMax (Thermo-Fisher) for polyI:C transfections and with Lipofectamine 2000 (Thermo-Fisher) for plasmid transfections. Cells were treated with indicated units of interferon-β for 16 hours.

### PolyA Luciferase plasmids

Firefly Luciferase with N terminal HA affinity tag was inserted with tandem poly A runs using the following forward primers and a stop codon containing reverse primer using the gateway cloning system.

4A_FLUC_BP_FOR GGGGACAACTTTGTACAAAAAAGTTGGC**AAAA**ATGTACCCATACGATG

8A_FLUC_BP_FOR GGGGACAACTTTGTACAAAAAAGTTGGC**AAAA AAAA**ATGTACCCATACGATG

12A_FLUC_BP_FOR GGGGACAACTTTGTACAAAAAAGTTGGC**AAAA AAAAAAAA**ATGTACCCATACGATG

20A_FLUC_BP_FOR GGGGACAACTTTGTACAAAAAAGTTGGC**AAAA AAAAAAAAAAAAAAAA**ATGTACCCATACGATG FLUC_BP_STOP_REV GGGGACAACTTTGTACAAGAAAGTTGGctacacggcgatctttccgcc

### Luciferase Assay

Cells were transfected with 1:1 ratio of plasmids encoding firefly luciferase (with varied polyA sequence lengths within the 5’UTR) and a control renilla luciferase plasmid using lipofectamine 2000. Luciferase activity was measured using a Dual-Glo reagent kit (Promega) and a Glomax (Promega) luminometer following manufacturer instructions.

### Virus infection, plaque assays

Vaccinia virus strain WR was a generous gift of R. Condit. For viral titering assays, cells were seeded in 24 well plates and grown overnight, followed by addition of 10,000 PFU/well vaccinia virus. Twenty-four hours after infection, resulting vaccinia virus was harvested by freeze/thaw lysis of infected cells. The resulting supernatant was serially 10-fold diluted in 24-well plates in DMEM + 10% FBS and overlaid on BSC40 cells at 80% confluency. 48 hours later, the media was aspirated, and the cells were stained with 0.1% Crystal Violet in 20% ethanol, and then destained with 20% ethanol. Initial viral conctractions were determined by manually counting plaques. For proteomic, RNAseq and western blot analyses, vaccinia virus was added at an MOI=5 and incubated for the indicated time before cells were harvested.

### RQC FACS assay

Cells were transfected with equal concentration of the Dual fluorescence reporter containing K20 internal repeat or K0 control in 293T cells. Fluorescence levels were measured using X-20 Fortessa (BD) instrument as done previously(Sundaramoorthy et al. 2017).

### RNA-Seq

Total RNA from mock or vaccinia infected cell lines were extracted using Trizol reagent (Invitrogen). Illumina Stranded Truseq total RNA kit was used for library preparation. Total RNA depeleted of Ribosomal RNA using Ribozero was converted to cDNA and single read 50 bp library was sequenced on Illumina Hi-Seq instrument (IGM core UCSD). The demultiplexed and de-indexed Fastq files were processed using galaxy (Afgan et al. 2018). Briefly, FASTq files were quality control checked using FASTQC (https://www.bioinformatics.babraham.ac.uk/projects/fastqc/) and aligned to the human genome (Hg38) using HISAT2/(Kim et al. 2015). The aligned BAM files were used for counting gene features using FEATURECOUNTS(Liao et al. 2014). DEBROWSER package (Kucukural et al. 2019) was used for differential gene expression analysis using DESEQ2(Love et al. 2014) and visualization of volcano plots and Heatmaps. FUNRICH (Pathan et al. 2015) was used for generating gene set venn diagrams and to identify overlapping gene sets.. To map reads to the vaccinia genome the FASTq files were aligned with custom indexed Vaccina WR genome (AY243312.1) using BWA(Li and Durbin 2009). HTseq counts(Anders et al. 2015) was used to identify mapped reads with genomic features. DEbrowser was used as previously indicated for identifying differentially expressed vaccinia genes. Heatmaps were generated using Graphpad Prism (8.4.3).

### Immunoblotting

Total cell lysates were prepared as follows. Cells after treatment were washed with PBS and trypsinized, pelleted and stored in −80. Frozen cell pellets were lysed using urea lysis buffer ((8Murea, 50mMTris-Cl, pH 8.0, 75mMNaCl, 1mMNaF,1 mM NaV, 1 mM b-glycerophosphate, 25 mM NEM)). Lysates were quantified using BCA. 25ug total protein was used for resolving on 4-20% Tris-glycine SDS page gels (Biorad). Resolved proteins were blotted to PVDF membranes (Bio-rad Immun-Blot) using Bjerrum semi-dry transfer buffer (48mMTris Base, 39mMGlycine-free acid, 0.0375% SDS, 20% MeOH, pH 9.2) in Biorad turbo transfer apparatus at 30V constant for 30 mins. Blots were incubated with relevant primary and HRP-conjugated (Promega) secondary antibodies and were detected using Clarity ECL reagent (Biorad) on a BioRad chemidoc XRS+ system.

### Mass-spectrometry

Cells were grown in a media containing either light (K0) lysine or ^13^C_6_ ^15^N_2_-labeled (K8) lysine (Cambridge Isotopes) and were processed for mass spectrometry as described previously (Markmiller et al. 2019). Briefly, cells were lysed using 8M urea lysis buffer and lysates were quantified for protein content using the BCA assay. 20μg of total cell extracts each from heavy and light SILAC labels were combined and diluted to a final urea concentration of 1M and then digested overnight with LysC enzyme (Promega) at a 1:100 (enzyme:protein) ratio. The digests were reduced with 1mM DTT for 30 min and then alkylated with 5mM IAA in a dark for 30min. The digests were desalted using Stage-Tip method and analyzed by LC-MS/MS as described (Markmiller et al. 2019). The resultant RAW files were analyzed using Andromeda/MaxQuant (version 1.6.0.16) using the combined Uniprot reviewed only database for Homo sapiens (2017) and Uniprot reviewed and unreviewed database for Vaccinia virus (2017). The default parameters were used except the enzyme was selected as LysC and ‘match between the runs’ and ‘requantify’ options were enabled in the MaxQuant settings.

---

**Supplementary figure S1.**
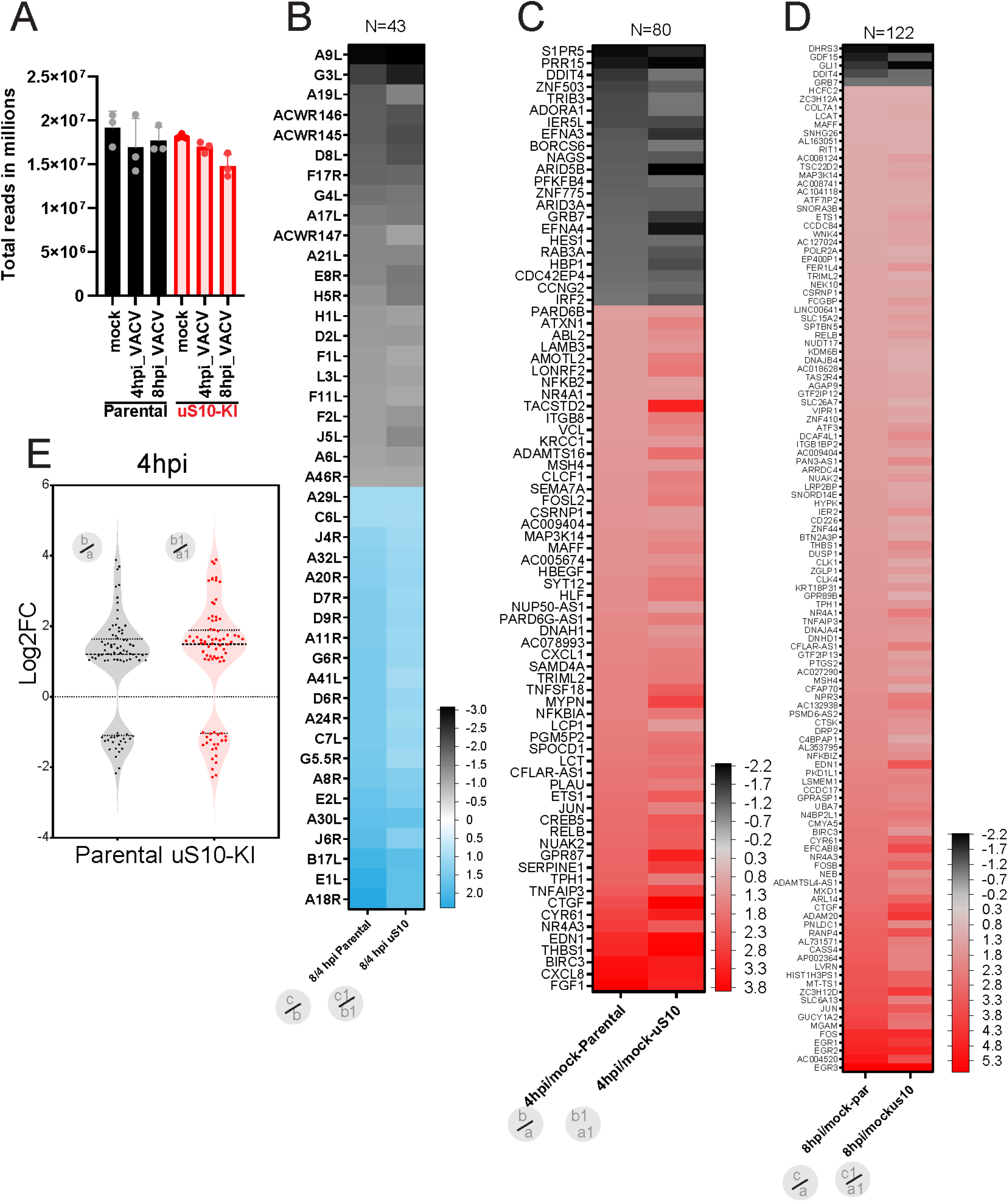
A) Bar-graph depicting the mean RNA-seq total reads in each indicated timepoints. Error bars denote SD of triplicate samples. B) Heat map depicting 43 differentially expressed (DE) vaccinia genes that overlap between the respective 8hpi contrasted with 4hpi within the same cell line. DE genes compared between parental and uS10-KI cell lines. Gene names are indicated on the left. Upregulated genes are represented in blue and downregulated in black. The log2 fold change range is between +2.3 to −3. C) Heat map depicting 80 differentially expressed human genes that overlap between parental and uS10-KI cell lines when comparing mock infected cells with those 4 hours after vaccinia virus infection. Upregulated genes are represented in red and downregulated in black. The log2 fold change range is between +3.8 to −2.2. D) Heat map depicting 122 differentially expressed human genes that overlap between parental and uS10-KI cell lines when comparing mock infected cells with those 8 hours after vaccinia virus infection. Upregulated genes are represented in red and downregulated in black. The log2 fold change range is between +5.6 to −2.2. E) Violin plot showing frequency distribution of the log2 ratios of the 80 common differentially expressed host genes comparing uninfected cells to those 4 hours post infection (hpi) within the indicated cell lines.

